# Vcflib and tools for processing the VCF variant call format

**DOI:** 10.1101/2021.05.21.445151

**Authors:** Erik Garrison, Zev N. Kronenberg, Eric T. Dawson, Brent S. Pedersen, Pjotr Prins

## Abstract

Since its introduction in 2011 the variant call format (VCF) has been widely adopted for processing DNA and RNA variants in practically all population studies — as well as in somatic and germline mutation studies. VCF can present single nucleotide variants, multi-nucleotide variants, insertions and deletions, and simple structural variants called against a reference genome. Here we present over 125 useful and much used free and open source software tools and libraries, part of `vcflib` tools and `bio-vcf`. We also highlight `cyvcf2`, `hts-nim` and `slivar` tools. Application is typically in the comparison, filtering, normalisation, smoothing, annotation, statistics, visualisation and exporting of variants. Our tools run daily and invisibly in pipelines and countless shell scripts. Our tools are part of a wider bioinformatics ecosystem and we consider it very important to make these tools available as free and open source software to all bioinformaticians so they can be deployed through software distributions, such as Debian, GNU Guix and Bioconda. `vcflib`, for example, was installed over 40,000 times and `bio-vcf` was installed over 15,000 times through Bioconda by December 2020. We shortly discuss the design of VCF, lessons learnt, and how we can address more complex variation that can not easily be represented by the VCF format. All source code is published under free and open source software licenses and can be downloaded and installed from https://github.com/vcflib.

**Author summary:** Most bioinformatics workflows deal with DNA/RNA variations that are typically represented in the variant call format (VCF) — a file format that describes mutations (SNP and MNP), insertions and deletions (INDEL) against a reference genome. Here we present a wide range of free and open source software tools that are used in biomedical sequencing workflows around the world today.

## Introduction

From its introduction in 2011 the VCF variant call format has become pervasive in bioinformatics sequencing workflows [1–3]. VCF is one of the important file formats in bioinformatics workflows because of its critical role in describing variants that come out of sequencing of DNA and RNA. VCF can describe single- and multi-nucleotide polymorphisms (SNP & MNP), insertions and deletions (INDEL), and simple structural variants (SV) against a reference genome [1]. Practically all important variant callers, such as GATK [4] and freebayes [5], produce files in the VCF format. The VCF file format is used in population studies as well as somatic mutation and germline mutation studies. In this paper we discuss the tools we wrote to process VCF and we shortly discuss strengths and shortcomings of the VCF format. We discuss how we can improve future variant calling in its contribution to population genetics.

An important part of the success of VCF that it is a relatively simple and flexible standard that is easy to read, understand and parse. This ‘feature’ has resulted in wide adoption by bioinformatics software developers. VCF typically scales well in bioinformatics workflows because files can be indexed [6], compressed [1, 7, 8] and trivially parallelized in workflows by splitting files and processing them independently, e.g. [5].

Here we present and discuss important tools and libraries for processing VCF in sequencing workflows: `vcflib`, `bio-vcf`, `cyvcf2`, `slivar` and `hts-nim`. These tools were created by the authors for the demands of large VCF file processing and data analysis following the Unix philosophy of small utilities as explained in the ‘small tools manifesto’ [9]. Development of these tools was often driven by the need to transform VCF into other formats, to digest information, to address quality control, and to compute statistics. The `vcflib` toolkit contains both a library and collection of executable programs for transforming VCF files consisting of over 30,000 lines of source code written in the C++ programming language for performance. `vcflib` also comes with a toolkit for population genetics: the Genotype Phenotype Association Toolkit (GPAT). Even though `vcflib` was not previously published, Google scholar shows 132 citations for 2020 alone — pointing to the github source code repository. Next to `vcflib`, we present `bio-vcf`, a parallelized domain-specific language (DSL) for convenient querying and transforming VCF; and we discuss `cyvcf2` [10], `slivar` [11], and `hts-nim` [12] as useful tools and libraries for VCF processing.

## Tools and libraries

### 2.1 vcflib C++ tools and libraries

At its core, `vcflib` provides C++ tools and a library application programmers interface (API) to plain text and compressed VCF files. A collection of 83 command line utilities is provided, as well as 44 scripts (Table 1 lists a selection). Most of these tools are designed to be strung together: piping the output of one program into the next, thereby preventing the creation of intermediate files, parallelize processing, and reducing the number IO operations. For example, the `vcflib vcfjoincalls` script includes the following pipeline (where `vt` is a variant normalization tool [13]; see Table 1 for the other individual steps by vcflib):

**Table 1.**
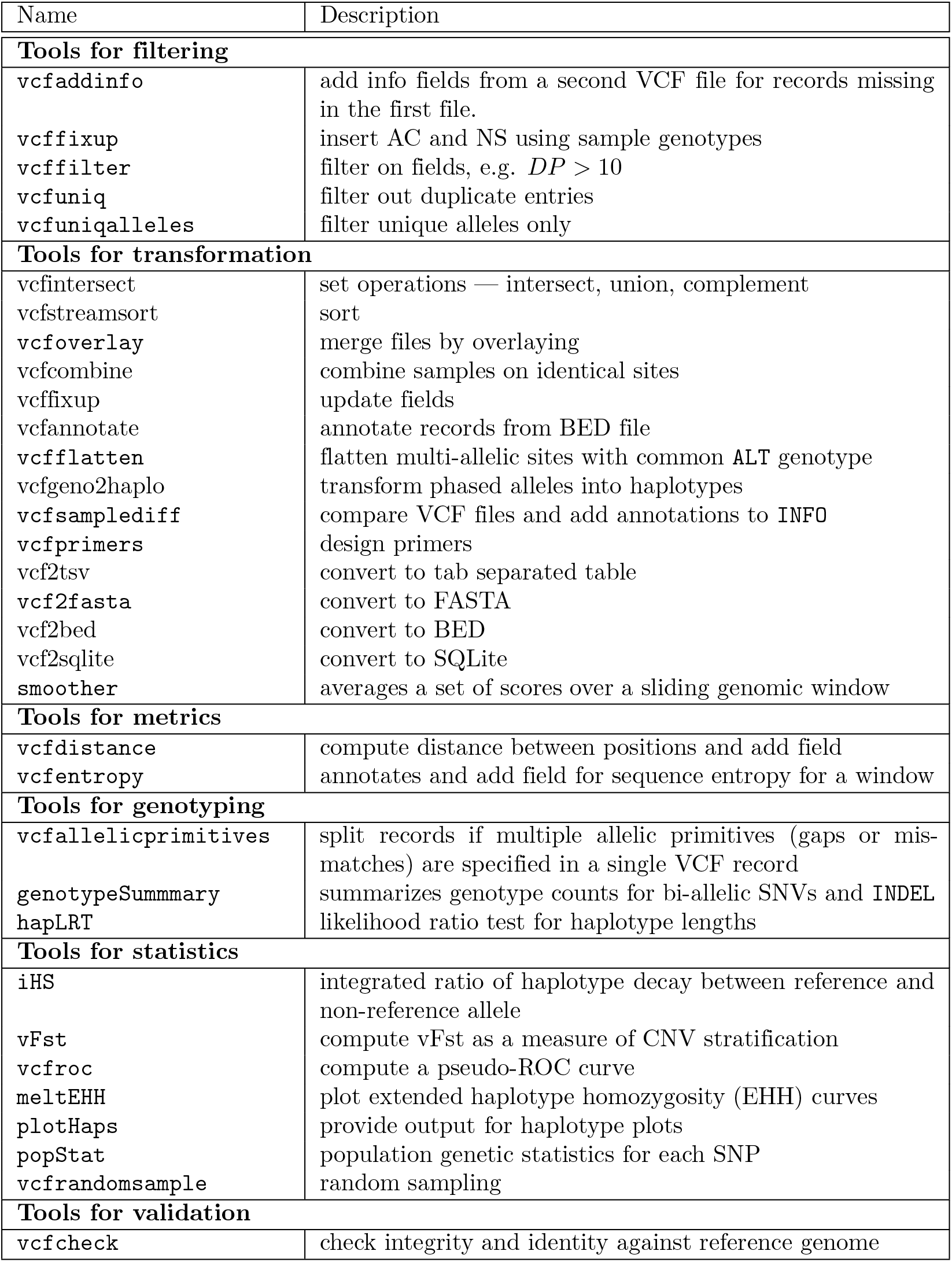
A selection of VCF processing tools included with `vcflib` (a full list of over 125 tools with descriptions and options can be found in the online `vcflib` documentation)

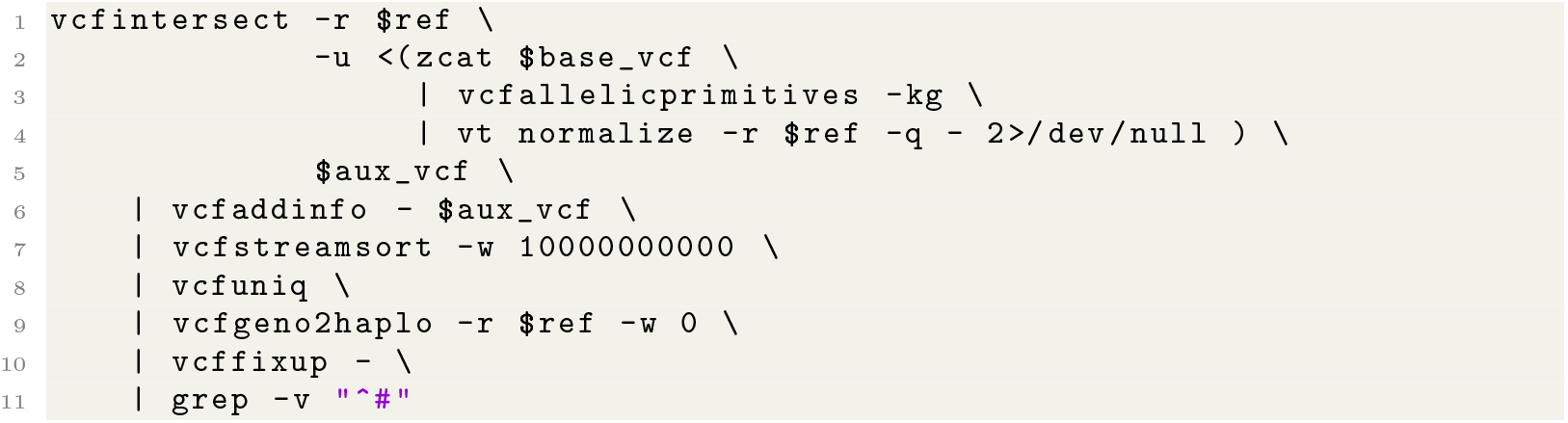

Such transformations and statistical functions provided in the toolkit were written for the requirements of projects such as the The 1,000 Genomes Project [14] and NIST’s genome in a bottle [15].

The VCF format is a textual file format: each line line describes a variant, i.e., a single nucleotide variant (SNV), an insertion, a deletion or a structural variant with rich annotation [1]. In a VCF line, fields are separated by the TAB character. Fields for chromosome, position, the reference sequence, the ALT alleles, and fields for quality, filter, INFO, FORMAT and calls for multiple samples are expected (see Fig. 1). To split fields, for example for ‘ALT T,CT’ another separator is used; in this case a comma. VCF makes use of many separators by splitting fields into subfields, subsubfields and so on: effectively projecting a ‘tree’ datastructure onto a single line. The advantage is that it is easy to view a VCF file and it is almost trivial to write a basic VCF parser and it is easy to add information to VCF, sometimes leading to unwieldy nested annotations. An evolving VCF ‘standard’ is tracked by the samtools/htslib project [17] and later amendments are particularly focused on more complex structural arrangements of DNA/RNA with ALT fields taking somewhat *ad hoc* creative forms, such as ‘A[3:67656[’ combined with an `INFO` field containing ‘SVTYPE=BND’ meaning that starting at reference position on a different chromosome, an ALT A nucleotide is followed by the sequence starting at chromosome 3 and position 67656. These SV annotations do away with some of the original simplicity of VCF. There are many such exensions introduced since the first publication of the VCF ‘standard’ that are used by specific SV tools and largely ignored by most VCF processing tools.

**Fig 1.**
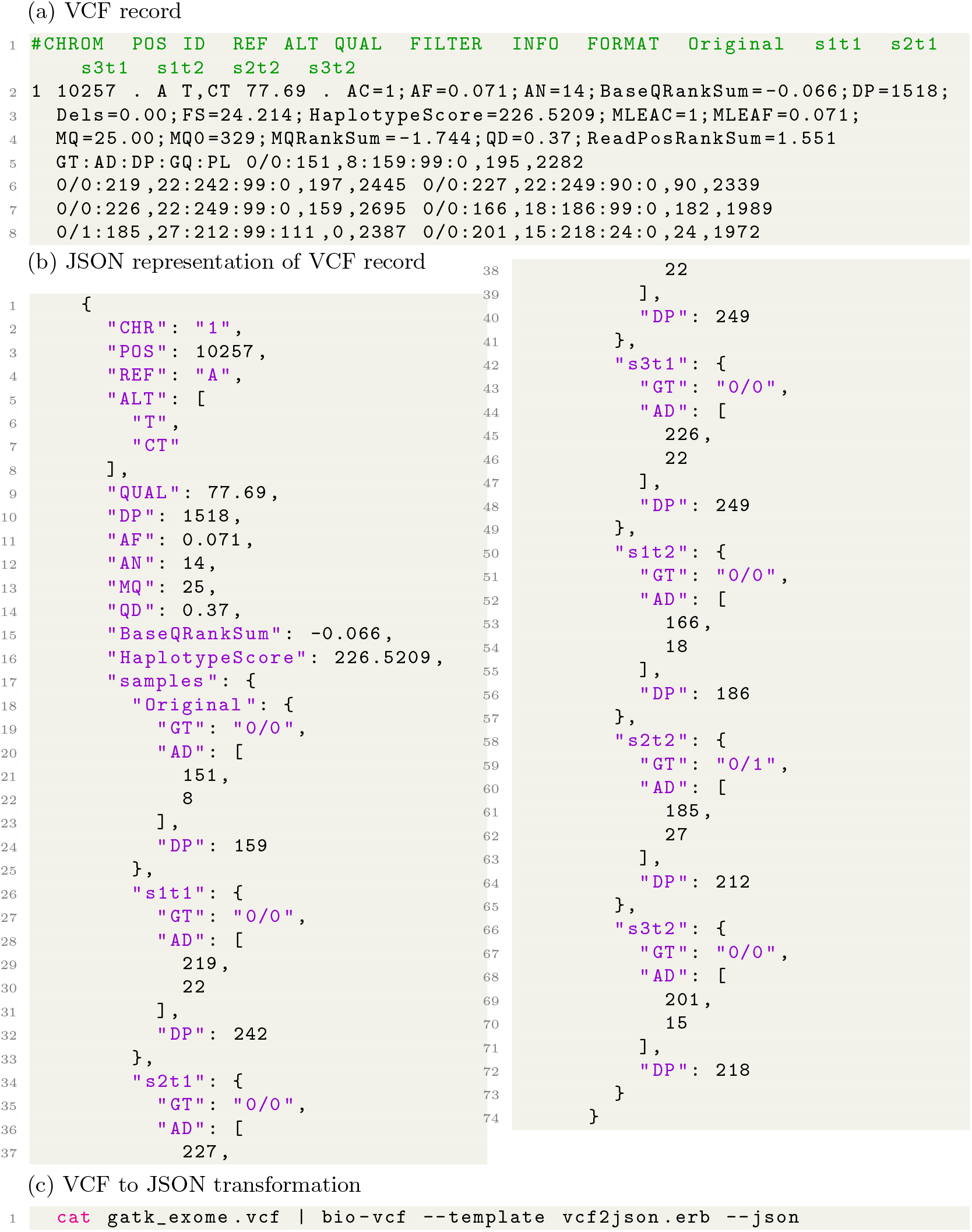
**Example of the VCF format and a VCF transformation to Javascript Object Notation (JSON) using `bio-vcf`** — (a) the line-based VCF record makes use of separators to split tab-delimited fields into subfields. Subfields are split with characters, =:; */* and so on. This splitting effectively projects a ‘tree-like’ datastructure that can also be represented as (b) a JSON record. JSON is used as a common data exchange format for databases and web-services. This example was generated with (c) the `bio-vcf` tool using a template [16]. `bio-vcf` transform data to any textual format, including RDF, HTML, XML etc. See also the `bio-vcf` section

The `vcflib` API describes a class `vcflib::VariantCallFile` to manage the reading of VCF files, and `vcflib::Variant` to describe the information contained in a single VCF record. The API provides iterators that are used in every included tool. For every record the tree-type hierarchy (Fig. 1) of information can be navigated in the record through interfaces to the fixed fields (CHROM, POS, ID, QUAL, FILTER, INFO) and sample-related fields (FORMAT, and samples). `vcflib` implements functions for accessing and modifying data in these fields; interpreting the alleles and genotypes in record; filtering sites, alleles, and genotypes via a domain-specific filtering boolean language; and reading and writing VCF streams.

In addition to the tools listed in Table 1, `vcflib` includes tools for genotype detection. For example, `abba-baba` calculates the tree pattern for four indviduals with an ancestral reference. `vcflib` includes a wide range of tools for transformation, e.g. `vcf2dag` modifies the VCF file so that homozygous regions are included as ‘`REF/.`’ calls. For each `REF` and `ALT` allele we can assign an index. These steps enable use of the VCF as a partially ordered directed acyclic graph (DAG). `vcfannotate` will intersect the records in the VCF file with targets provided in a BED file and `vcfannotategenotypes` annotates genotypes in the first file with genotypes in the second. `vcfclassify` generates a new VCF where each variant is tagged by allele class: `SNP`, `Ts/Tv`, `INDEL`, and `MNP`. `vcfglxgt` sets genotypes using the maximum genotype likelihood for each sample. `vcfinfosummarize` and `vcfsample2info` edit annotations given in the per-sample fields and adds the mean, median, min, or max to the site-level INFO. `vcfleftalign` Left-align INDELs and complex variants in the input using a pairwise REF/ALT alignment followed by a heuristic, iterative left realignment process that shifts INDEL representations to their absolute leftmost (5’) extent.

`vcflib` includes tools for phenotype association based on VCF files. We have developed a flexible and robust genotype-phenotype software library designed specifically for large and noisy NGS datasets. Wright’s F-statistics, and especially `Fst`, provide important insights into the evolutionary processes that influence the structure of genetic variation within and among populations, and they are among the most widely used descriptive statistics in population and evolutionary genetics (Fig 2). `Fst` is defined as the correlation between randomly sampled gametes relative to the total drawn from the same subpopulation [19]. `wcFst` is Weir & Cockerham’s `Fst` for two populations [20]. `pFst` is a probabilistic approach for detecting differences in allele frequencies between two populations and `bFst` is a Bayesian approach. `bFst` accounts for genotype uncertainty in the model using genotype likelihoods. For a more detailed description see [21]. The likelihood function has been modified to use genotype likelihoods provided by variant callers. There are five free parameters estimated in the model: each subpopulation’s allele frequency and `Fis` (fixation index, within each subpopulation), a free parameter for the total population’s allele frequency, and `Fst`. `pVst` calculates `Vst` to test the difference in copy numbers at each SV between two groups: *V st* = (*V t* − *V s*)*/V t*, where `Vt` is the overall variance of copy number and `Vs` the average variance within populations. `sequenceDiversity` calculates two popular metrics of haplotype diversity: *Pi* and extended haplotype homozygosity (`eHH`). *Pi* is calculated using the Nei and Li formulation [22]. `eHH` is a convenient way to think about haplotype diversity. When `eHH=0` all haplotypes in the window are unique and when `eHH=1` all haplotypes in the window are identical. The `vcfremap` tool attempts to realign, for each alternate allele, against the reference genome with a lowered gap open penalty and adjusts the CIGAR and `REF`/`ALT` alleles accordingly. These traditional and novel population genetic methods are implemented in the Genotype Phenotype Association Toolkit (GPAT++), part of the `vcflib` API. For example, permuteGPAT++, adds empirical p-values to a GPAT++ score. And `vcfld` computes linkage disequilibrium (LD). GPAT++ includes basic population stats (`Af`, `Pi`, `eHH`, `oHet`, `genotypeCounts`) and several flavors of `Fst` and tools for linkage, association testing (genotypic and pooled data), haplotype methods (hapLrt), smoothing, permutation, and plotting.

**Fig 2.**
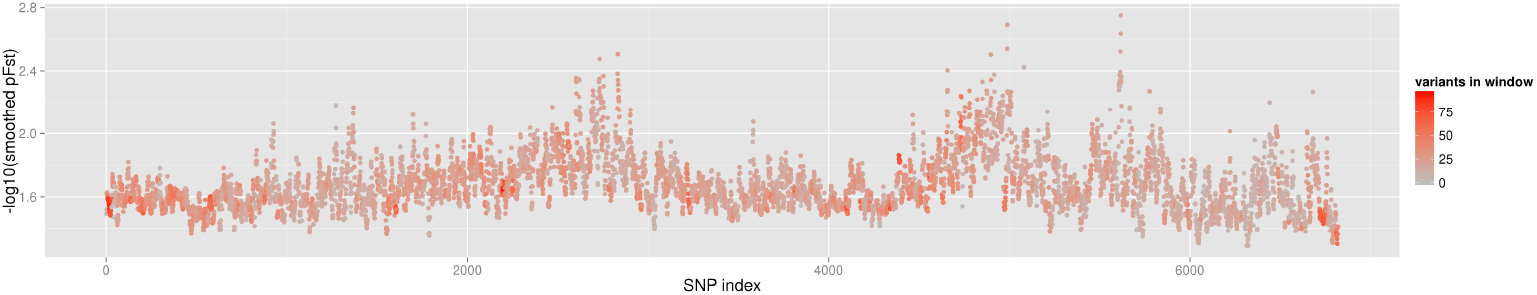
**Smoothed `pFst` (−*log*10) statistic with color coded number of variants in a window** — as computed by vcflib’s `pFst` and `smoother` tools [18].

`vcflib` includes tools for genotype statisics. `vcfgenosummarize` adds summary statistics to each record summarizing qualities reported in called genotypes. It uses `RO` (reference observation count), `QR` (quality sum reference observations) `AO` (alternate observation count), `QA` (quality sum alternate observations). The `normalizeHS` is used for `iHS` and `XP-EHH` scores [23].

A full list of over 125 commands and functionality can be found on the website, as well as documentation and examples of application [18].

### 2.2 Bio-vcf and Slivar flexible command-line DSL filters and transformers

#### 2.2.1 bio-vcf

Compared to `vcflib` with its many dedicated command line tools, `bio-vcf` takes a different approach by providing a single command line tool that uses a domain specific language (DSL) for processing the VCF format. Thanks to a dynamic interpretation of the VCF tree representation (see Fig. 1) all data elements in a VCF header or record can be reached using field names and their sub names. For example the following is a valid select filter:

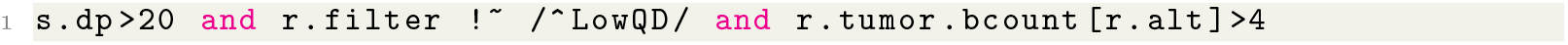

which selects all variants where the sample depth field `s.db` is larger than 20, where the `FILTER` field of a record `r` does not start with the letters *LowQD* (note it uses a Perl/Ruby-style regular expression or regex [24]), and where the tumor bcount of the *ALT* allele is larger than 4. The letter ‘r’ represents a record or line in a VCF file and the letter ‘s’ stands for each sample in a record.

The naming of variables, such as s.dp and r.tumor.bcount, is inferred from the VCF file itself, so if a VCF has different naming conventions they are picked up automatically.

`bio-vcf` typically reads from the terminal STDIN and writes to STDOUT. The following full command line invocation reads VCF files and filters for chromosomes 1–9 where the quality (r.qual) is larger than 50. It also checks for non-empty samples where the sample read depth is larger than 20. For each selected record with `--eval` it outputs a BED record (the default output is the VCF record itself, useful for filtering):

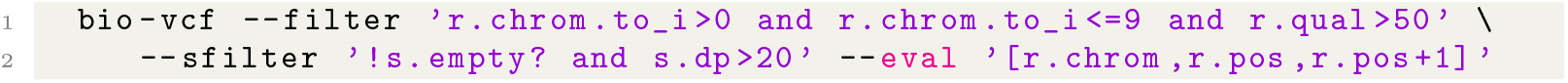

For comparisons and for output, fields can be converted to integers, floats and strings with to i, to f and to s respectively. Note that these are Ruby functions and, in fact, all such Ruby functionality is available in `bio-vcf` statements. For extreme flexibility `bio-vcf` even supports *lambdas* which makes for very powerful queries and transformations. For example, to output the count of valid genotype fields in samples one could use

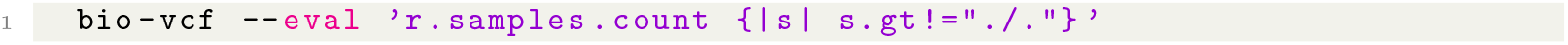

where `count` is a function that invokes the *lambda* `s.gt!=“./.”`, i.e., where the genotype `gt` of sample `s` is not equal to “./.”. Sample ‘`s`’ is passed as a parameter.

Because of the flexibility of `bio-vcf` almost all imaginable data queries can be executed. `bio-vcf` was implemented for processing large VCF files and is fast because it is designed make use of multi-core processors (using Linux parallelized copy-on-write, i.e. a technique where RAM is shared between processes). `bio-vcf` is also ‘lazy’ which means that it only parses fields when they are used. For example, in the above query, only the sample *GT* field is unpacked and parsed to get a result. All other data in the record is ignored by the query and not evaluated. This contrasts largely with most VCF parsers in use today.

Finally, `bio-vcf` comes with a full parser and lexer that can tokenize the VCF file header and transform that in some other format. For example, the command

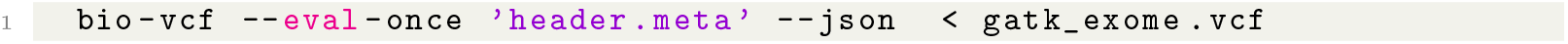

will turn the metadata information passed by GATK [4] into a JSON document. To get a full JSON document of the VCF file use a template that looks like

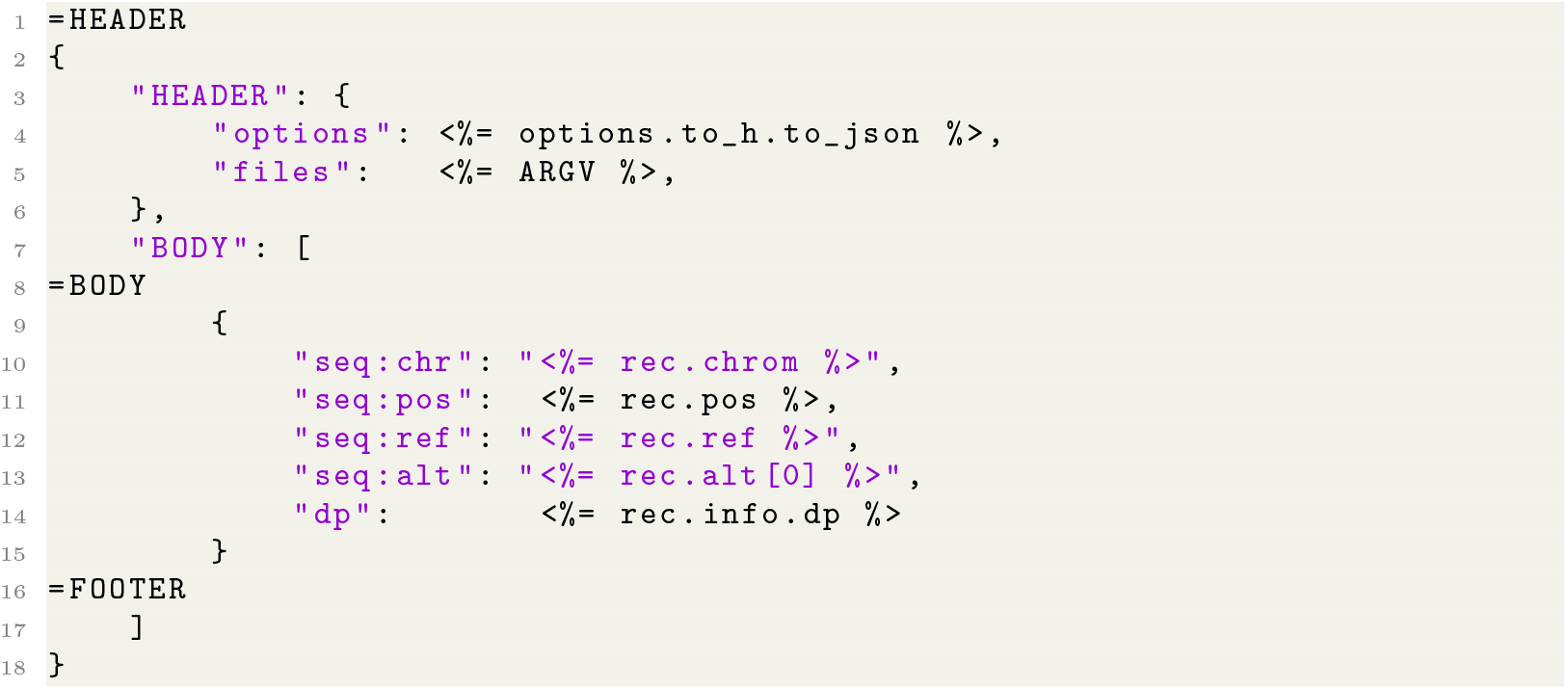

and run

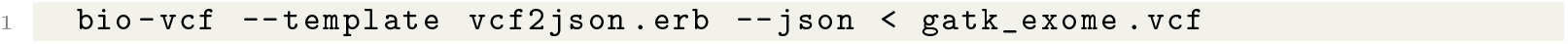

The high expressiveness and adaptable parsing makes `bio-vcf` a very powerful tool for searching, filtering and rewriting VCF files. See the `bio-vcf` website for full information on record and sample inclusion/exclusion filters, generators, multicore performance, field computations, statistics, genotype processing, set analysis and templates for user definable output, including templates for output of VCF header information and records for RDF, JSON, LaTeX, HTML and BED formats [16].

#### 2.2.2 Slivar

Similar to `bio-vcf`, `slivar` allows users to specify simple expressions for filtering and annotation [11]. Whereas `bio-vcf` uses Ruby to supply the DSL, `slivar` uses Javascript. `slivar` has built-in pedigree support for the samples so that, for example, a single expression can be applied to every trio (mother, father, child) to identify *de novo* variants:

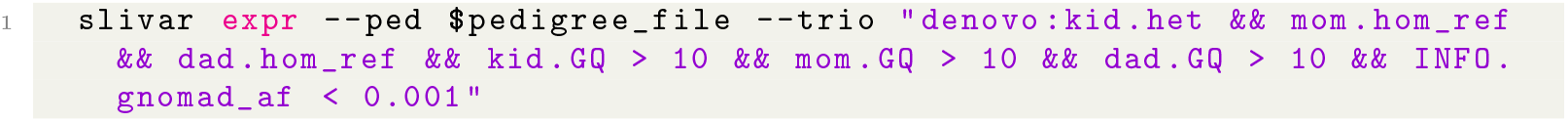

The expression above checks the genotype pattern along with genotype quality and limits to rare variants by `INFO.gnomad af`*<* 0.001.

Expressions on families (including multigenerational) and arbitrary groups are supported so that, for example, expressions can be applied to tumor-normal pairs using `tumor` and `normal` labels.

### 2.3 VCF programming libraries

VCF programming libraries are mainly useful when direct calls to `vcflib` and `bio-vcf` command line tools proves too limited. The Bio* libraries, e.g., biopython [25], bioperl [26], bioruby [27] and R’s CRAN [28], contain VCF parsers that may be useful. But a first point of call may be `vcflib` itself as it is also a C++ programming library and in addition to being an integral part of the `vcflib` tools mentioned here is used by, for example, the freebayes variant caller [5].

Of particular interest is the fast `cyvcf2` library that was started in 2016 with htslib [17] bindings and is actively maintained by co-author Pedersen today [10]. Similar to `bio-vcf` it presents a DSL-type language that can be used in Python programming. Meanwhile hts-nim [12] contains bindings for the Nim programming language with similar syntax and functionality. For example a nim bin counter (part of hts-nim-tools/vcf-check [12]) can be written as:

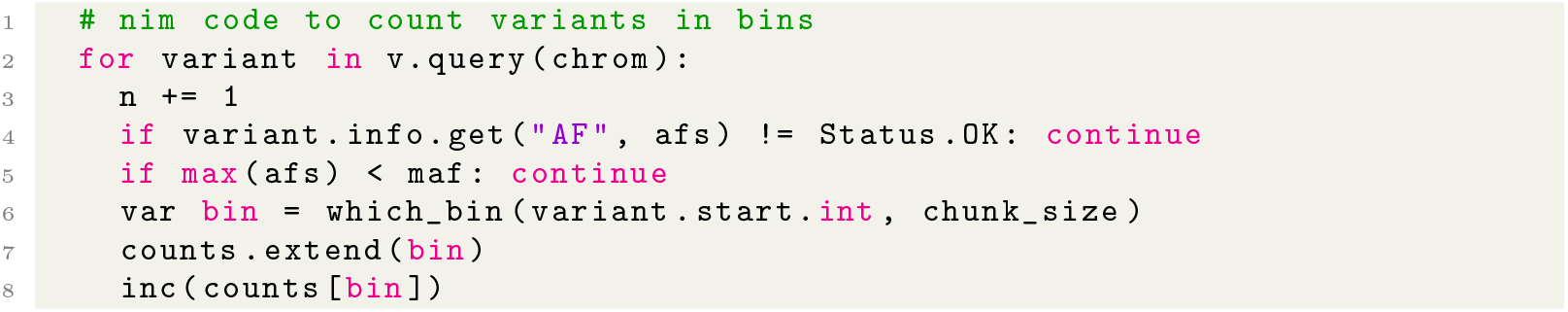

Nim is a statically typed compiled language that looks very similar to Python and, because it transpiles to C, Nim has a much faster runtime and can link without overheads against C libraries such as `vcflib` and `htslib`.

Finally, another tool of interest, by the same author, is `vcfanno`; written in the Go language and allows annotations of a VCF with any number of INFO fields from any number of VCFs or BED files. `vcfanno` uses a simple conf file to allow the user to specify the source annotation files and fields and how they will be added to the info of the query VCF [29].

## Discussion

Ten years is a long time in bioinformatics and the VCF file format is starting to show its age. Not only is the VCF format redundant and bloated with duplication of data, a more important concern is that the VCF format does not accommodate interesting complex genomic variations, such as complex and nested variants, such as superbubbles, ultrabubbles, and cacti [30–32]. An even more important shortcoming of VCF is that it always depends on a single reference genome, resulting in variant calling bias and missing out on variation not represented in the reference [32]. One solution is to work with multiple reference genomes, but comparing VCF files from different reference genomes is challenging — even for different versions of one reference genome.

To address such challenges the authors are actively working on pangenome approaches that store variation in a pangenome graph format, e.g. [32–37]. Pangenomes can incorporate multiple individuals and multiple reference genomes. Pangenomes can cater for very complex structural variation. Pangenomes are also efficient in storing information, including metadata, without redundancy. In effect, pangenomes cater for a ‘lossless’ view of all data at the population level. This largely differs from VCF-type data because, despite mentioned data size, a generated VCF implies a data reduction step — or data loss — that effectively disconnects variants from each other and related features, such as quality metrics. This means that rebuilding the original data from VCF files is virtually impossible. In contrast, with the newer pangenome formats it is possible to rebuild sequences independent of the underlying complexity of features. Having a full view of the data makes downstream analysis, such as population genotyping, more powerful with improved results, e.g. [32].

The VCF file format has become a crucial part of almost all sequencing workflows today. The design and presentation of the VCF file format can set the norm for designing future file formats [2, 3], but we can also learn from its mistakes. In this paper we wrote VCF ‘standard’ consistently between quotes because, even though there exists a standardization effort — now at VCFv4.3 [3] — VCF is flexible by design, alternative VCF standards are introduced (e.g. [38]) and most tools take liberties when it comes to producing VCF files. Therefore all VCF parsers have to take a flexible approach towards digesting input data and ignore input data that is not understood.

We recognise that the success of a file format requires a crucial focus on having an early ‘standard’ that is both easy to understand and flexible enough to grow, in line with the success of other bioinformatics file formats, such as SAM/BAM [3] and GFF [39]. Biology is a fast moving field and it is impossible to predict how a file format is going to be used in a (near) future. The downside of such flexibility is that older software may not support features that were added later. One of the weakest aspects of the VCF format is its metadata: next to *ad hoc* metadata in records (see Fig.1), the header record requires specialized parsing and ignores existing ontologies.

Also for the VCF records, robust validation, error checking and correctness checking is virtually impossible. Great attention should therefore be paid to any amendments to an earlier standard to keep backward compatibility when possible. VCF and many other formats in bioinformatics use layered character separators as a grammar for defining a tree structure of data (see Fig. 1). This type of format requires specialized *ad hoc* parsers for every format. In the future, when designing new formats, we strongly suggest to base a new format on existing standards such as JSON, JSON-LD and RDF web formats for storing hierarchical data and graph data respectively. Each of these formats has efficient storage implementations. A future format should also benefit from reusing existing ontologies or create and champion a new ontology, if one does not exist, so data becomes easily shareable, comparable and queryable and living upto FAIR requirements [40]. Not only are JSON, JSON-LD and RDF natively and efficiently supported by most computer languages, they are also more easily embedded in existing infrastructure, such as NoSQL databases.

### Software development and distribution practices

In this paper we present three types of tools that mirror three common approaches in bioinformatics towards large data parsing. First are `vcflib` Unix style command line tools where each tool does a small job [9]. Second are `bio-vcf` and `slivar`-style extremely flexible command line DSLs. And third are programming against libraries that can be called from programming languages, such as `cyvcf2` and `hts-nim`, as well as `vcflib` and `bio-vcf`.

A wide range of solutions exist for VCF processing that make use of these three approaches and functional overlap is found between `vcflib`, `bio-vcf`, `cyvcf2`, the original vcftools [2], bcftools [7] and the existing Bio* programming libraries, such as biopython [25], bioruby [27] and biojava [41]. vcftools and bcftools provide annotation, merging, normalization and filtering capabilities that complement functionality and can be combined in workflows with `vcflib` and `bio-vcf`. These solutions together provide a comprehensive and scalable way of dealing with VCF data and every single tool represents a significant investment in research and software development. Therefore, before writing a new parser from scratch, we strongly suggest to first study the existing solutions. In the rare case a new tool is required it may be an idea to merge that with existing projects so everyone can benefit.

Once software is written, it is important software development and maintenance continues. In the biomedical sciences it is a clear risk for projects to get abandoned once the original author moves on to another job or other interests; partly due to a lack of scientific recognition, attribution and reward [42]. We note that with the `pyvcf` project, for example, this has happened twice and the github contribution tracker shows no more contributions by a project owner. This means no one is merging changes back into the main code repository and the code is essentially unmaintained. `vcflib`, `bio-vcf` and `cyvcf2`, in contrast, show a continuous adoption of code contributions thanks to the original authors encouraging others to take ownership and even release versions of the software. We also recognise the importance of creating small tools that can interact with each other following the Unix philosophy.

For overall adoption of software solutions it is important the tools and documentation get packaged by software distributions, such as Bioconda [43], Debian [44] and GNU Guix [45, 46]. Bioconda downloads are a good estimation of relative popularity because they tend to represent actual installations. `vcflib`, for example, was installed over 40,000 times and `bio-vcf` was installed over 15,000 times through Bioconda by December 2020. `vcflib` is also an integral part of the freebayes variant caller with an additional 110,000 downloads through Bioconda.

### Future work

Software development never stands still. With new requirements tools get updated. With the evolving VCF ‘standard’ and associated tooling for pangenomes and reference based approaches we keep updating our tools and libraries. We intend to add more documentation and regression tests. One recent innovation in `vcflib` is the generation of Unix ‘man’ pages from markdown pages and the `--help` output from every individual tool and script that also doubles as regression tests. This documentation is always up to date because when it goes out of sync the embedded tests fail. We think it will also be useful to add tool descriptions in the common workflow language (CWL) format [46–48]. CWL definitions allow easy sharing of tool components between sequencing workflows. The scenario will be to write a CWL tool definition that can be converted to documentation and running tests.

Despite our criticism of VCF, VCF as a file format is likely to remain in use. To replace VCF most existing tools and workflows used in sequencing would have to be rewritten. Pangenome tools, in principle, are capable of producing reference guided VCF files from GFA graph structures. These tools guarantee compatibility with upstream and downstream analysis workflows. We predict that pangenome approaches will play an increasingly important role in sequence analysis and, at the same time, VCF processing tools will remain in sequencing workflows for the forseeable future.

## Acknowledgments

These free software tools are a community effort and benefit from ideas by hundreds of people. We thank all collaborators, supervisors and other contributors. We particularly want to thank software packagers, such as Andreas Tille and Michael R. Crusoe for providing these tools in Debian. We also want to thank Brad Chapman and Torsten Seemann for the Bioconda packages and Efraim Flashner for the GNU Guix packages.

